# Brain Decoding from a Deep Neural Network of Visual Memory Reveals the Maturation Trajectory of Memory Systems across Ages 3-7

**DOI:** 10.64898/2026.11.03.763814

**Authors:** Xiaohan (Hannah) Guo, Yuan Chang Leong, Wilma A. Bainbridge

**Author notes:** Correspondence should be directed to Xiaohan (Hannah) Guo, Department of Psychology, University of Chicago, 940 East 57th Street, Chicago, IL, 60637,.

## Abstract

Memories from the first four years of life are largely inaccessible by adulthood, yet these early experiences profoundly shape cognition throughout development. Understanding this phenomenon of infantile amnesia is critical for revealing the mechanisms of early memory development. Here, we developed a new content-based behavioral measure of memory that quantifies how closely children’s memory patterns resemble those of adults, which we term the *memory maturation score*. Using a movie-watching fMRI dataset spanning age 3 to adulthood, we trained a cross-validated model on adults’ movie-evoked brain activity to predict adult’s propensity to remember each movie snippet, estimated by ResMem, a deep neural network trained to predict visual memorability. We then applied the model to each child’s brain activity while viewing the same movie to assess their memory maturation (i.e., cross-decodability from adult patterns). Children exhibited adult-like neural patterns of memory around age 4, extending prior behavioral findings. Structural analyses showed that the volume of the left posterior hippocampus, but not age or total brain volume, significantly predicted children’s memory maturation. Memory maturation around age 4 thus likely reflects age-related neural development of hippocampal memory systems, rather than an age increase itself. Functional searchlight analyses revealed a trajectory of memory maturation: the inferior temporal gyrus reflected mature memory-related processing from age 3 onward, the left posterior hippocampus by age 5, and higher-order regions involved in social-affective processing and multimodal integration by age 7. Together, these findings suggest that hippocampal maturation between ages 3 and 5 drives the maturation of memory patterns.

**Significance Statement:** Despite decades of research establishing potential mechanisms of infantile amnesia, little is known about the maturation of neural memory representations across development, limiting progress in developmental memory research and educational applications. This study reveals a developmental trajectory of memory: children show adult-like processing in the IT area by age 3, the hippocampus by age 5, and higher-order regions by age 7, reflecting mature perceptual, memory, and multimodal and social-affective processing, respectively. The left posterior hippocampus is crucial for the maturation of adult-like memory patterns. These findings show that (i) ResMem DNN can help educators select dynamic stimuli that are memorable to children by age 4, and (ii) our cross-validated model enables assessment of memory maturation without explicit memory measures.

## Introduction

Can you remember the thrill on your parents’ faces when you first smiled, spoke your first word, or took your first step? For most people, the answer is no. No matter how much we wish to hold onto those warm, early moments of our lives, they remain beyond our reach. Perhaps some moments are more difficult for the developing brain to retain, while others leave lasting traces that shape learning and development. Although these early memories have largely faded by adulthood, they continue to shape who we become. Recovering insight into this inaccessible period of life could deepen our understanding of how early experiences shape later cognition. Building on advances in computational methods, the present study adopts a content-specific perspective to examine how children process naturalistic stimuli across development, with a particular focus on the neural maturation underlying developmental changes in perceptual and memory processing. In doing so, we aim to provide a more mechanistic understanding of infantile amnesia in humans.

Infantile amnesia is a robust, cross-cultural phenomenon whereby most adults have no conscious memories from their first three to four years of life (1). This transitions into childhood amnesia, a related phase characterized by a scarcity of retrievable memories from ages five to seven. Such a lack of early memories cannot be fully explained by memory decay (i.e., forgetting; 2, 3) or interference (i.e., newer memories overwriting or competing with older traces; 4) alone, motivating investigation into what determines which information children tend to remember. Research on infantile amnesia has primarily focused on individual-specific capacity measures to quantify the development of memory, such as the number of trained events produced by children (5, 6) or the number of autobiographical episodes recalled by children (3, 7), while largely overlooking item-specific memories within episodes that fade during early childhood. To our knowledge, only one prior study has examined specific memory contents of children, albeit at a relatively coarse level. DeBoer and colleagues (8) found that children showed better recall in late childhood for events that had been described with higher quality at age 3, as indexed by qualitative ratings such as structural coherence. However, simply knowing that children retain more memories as they age, and that better-encoded memories are better retained later in life, does not tell us which aspects of information are more likely to be remembered or why across development. This lack of understanding of what information tends to be remembered at each moment in an event across children limits our understanding of what and how memories are retained as infantile amnesia fades. This highlights the need to study infantile amnesia from a content-specific perspective by identifying the types of information that are preferentially encoded during early childhood.

One tractable way to examine the contents encoded at the cusp of infantile amnesia is to compare the episodic visual information (e.g., images, video clips, etc.) that children remember with that remembered by adults. Prior work has shown that adults consistently remember and forget the same visual content across people—including scenes (9), abstract symbols (10), abstract artworks (11), and dynamic visual-only and audiovisual monologues (12). These memory consistencies reflect shared sensitivity in adults’ visual memory systems to the semantic aspects of stimuli (13), rather than low-level visual contrast, aesthetics, and emotion (14, 15). This stimulus-intrinsic *memorability* accounts for more than half of the variance in adult memory performance (16) and can be predicted by ResMem, a deep neural network (DNN) trained to predict how likely an image is to be remembered across adults (17). ResMem therefore provides a proxy for mature adult memory patterns of visual content. Because the shared memory patterns can also be measured from brain data during perception (18), perceptual neuroimaging provides a particularly valuable approach for studying the memory of young children, for whom high-quality memory data can be difficult to collect.

These findings on memorability in adults suggest that children’s memory patterns can be studied by first identifying when their behavioral memory patterns resemble those of adults, and then examining the neural developmental trajectory that gives rise to these similarities. To compare behavioral memory patterns between adults and young children, prior research compared scene memorability scores from a paired-associative memory task in young children with ResMem-predicted memorability, which reflects adults’ mature memory patterns (19). Surprisingly, ResMem trained on adults’ memory data predicted young children’s memory for novel stimuli in a novel task paradigm. Critically, children exhibited mature behavioral memory patterns by age 4. Building on these behavioral findings, here we develop a computational approach that leverages ResMem-derived memorability patterns to quantify how closely children’s memory-related brain signatures resemble those of adults, yielding a *memory maturation score* per child. The prior behavioral finding highlights the temporal convergence between the maturation of children’s behavioral memory patterns and the attenuation of infantile amnesia, making ages 3 to 4 a critical developmental window for understanding what and how children remember as they transition out of infantile amnesia.

The maturation of memory patterns and the attenuation of infantile amnesia also share a common neural substrate—the hippocampus. Prior functional magnetic resonance imaging (fMRI) studies have shown that adults exhibit more similar brain activation patterns in the medial temporal lobe, including the left hippocampus, when viewing memorable versus forgettable images (20), suggesting that adult behavioral memory patterns are supported by hippocampal processing. Likewise, animal studies over the past decade have demonstrated a causal relationship between decreased hippocampal neurogenesis and the attenuation of infantile amnesia (21). Forgotten infantile fear-conditioning memories can also be reinstated in laboratory rats through optogenetic stimulation of the hippocampus (22), suggesting the role of the hippocampus in mitigating infantile amnesia. Recent fMRI studies in human infants have further illuminated that greater hippocampal activity during encoding predicts subsequent memory for images by age one (23), demonstrating the crucial role of the hippocampus in early memory encoding. Together, these findings indicate that infantile amnesia reflects developmental constraints on memory systems involving the hippocampus rather than a complete inability to encode or store experiences (1, 2). Nevertheless, infantile amnesia in human early childhood remains elusive. Many questions remain unanswered (see review: 24), including the developmental trajectories of the different mechanisms that support adult memory, and the properties of early memory traces during encoding. The present study provides a first step toward addressing these questions by comparing the memory brain signatures across early and middle childhood with those observed in adults.

Understanding children’s memory maturation has broad implications. For developmental researchers, our findings provide an objective tool for comparing memory maturation across children and studies without requiring explicit memory assessment. For the general public, understanding what children actually remember could inform educational and parenting practices, guide AI systems that model how early experiences are “forgotten,” and potentially support therapeutic approaches to early childhood trauma by identifying which aspects of early experiences may persist and influence later life, even in the absence of conscious memory.

Here, we use a content-based approach to reveal how children’s memories diverge from adults’ by computing a *memory maturation score* for each child using a cross-validated model (Fig. 1). To preview, we find that children begin to show brain signatures of adult-like memory patterns by around age 4. This maturation is predicted by the left posterior hippocampus, both functionally and structurally, rather than by general increases in age, gender, or brain size. Children show mature processing in visual-processing regions (inferotemporal gyrus) by age 3, memory-processing regions (left posterior hippocampus) by age 5, and social-affective and multimodal integration regions by age 7. This neural developmental trajectory reveals increasingly enriched memory representations during encoding across early childhood.

**Fig. 1.**
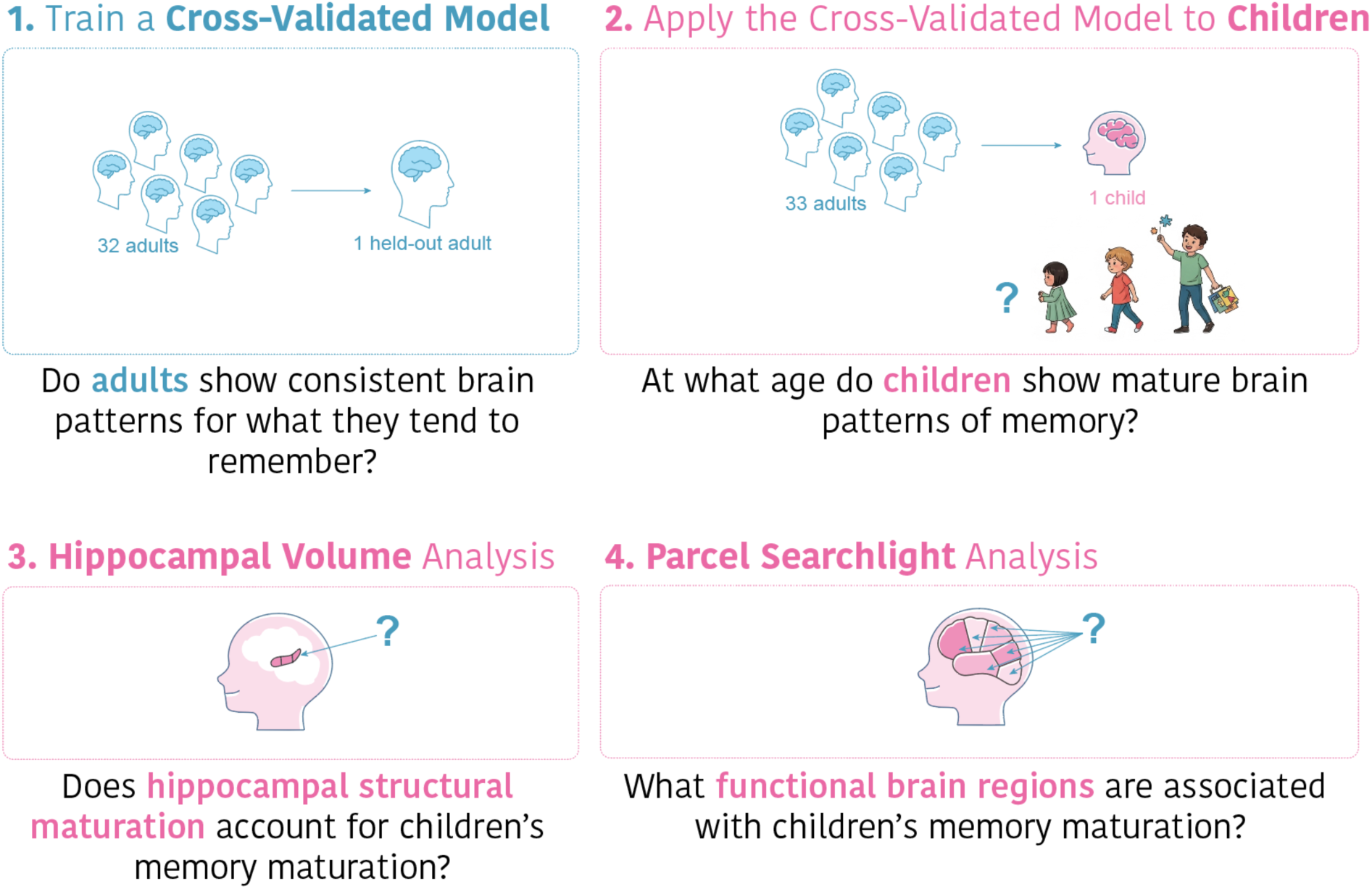
Overview of the Research Questions Addressed in This Study. **1.** To determine whether adults exhibit consistent neural patterns for information they tend to remember, we trained a leave-one-subject-out cross-validated model to predict movie memorability (estimated by the ResMem deep neural network) from adults’ whole-brain activity. **2.** To determine at what age children exhibit mature neural patterns of memory, we applied the adult-trained cross-validated model to each child between ages 3 and 10. A higher correlation between the model-predicted memorability from a child’s brain data and the ResMem-estimated movie memorability indicates greater memory maturation. **3.** To examine whether structural maturation of the hippocampus predicts children’s memory maturation, we fit linear regression models using hippocampal subfield volumes to predict each child’s memory maturation score (i.e., the prediction accuracy of the cross-validated model). **4.** To identify the functional brain regions associated with children’s memory maturation, we repeated the adult-to-child cross-validated analysis using brain activity from each parcellated brain region separately.

## Results

### The Movie-Watching fMRI Dataset

We used an existing fMRI dataset originally designed to study children’s theory of mind (*N_adult_*=33; *N_child_*=116, aged 3-10; 25). Participants watched a 5.6-min muted Disney Pixar movie, *Partly Cloudy* (24 frames/s), while undergoing whole-brain functional MRI (TR=2s, see Methods and Materials for more scan acquisition details). T1-weighted anatomical scans were also acquired. All functional data were preprocessed and registered to the Montreal Neurological Institute (MNI) template. We focused on the main movie segment (5.2 min, excluding the opening Pixar logo and closing credits) and the corresponding brain signatures (155 TRs), accounting for a 4-s hemodynamic lag (Fig. 2A).

**Fig. 2.**
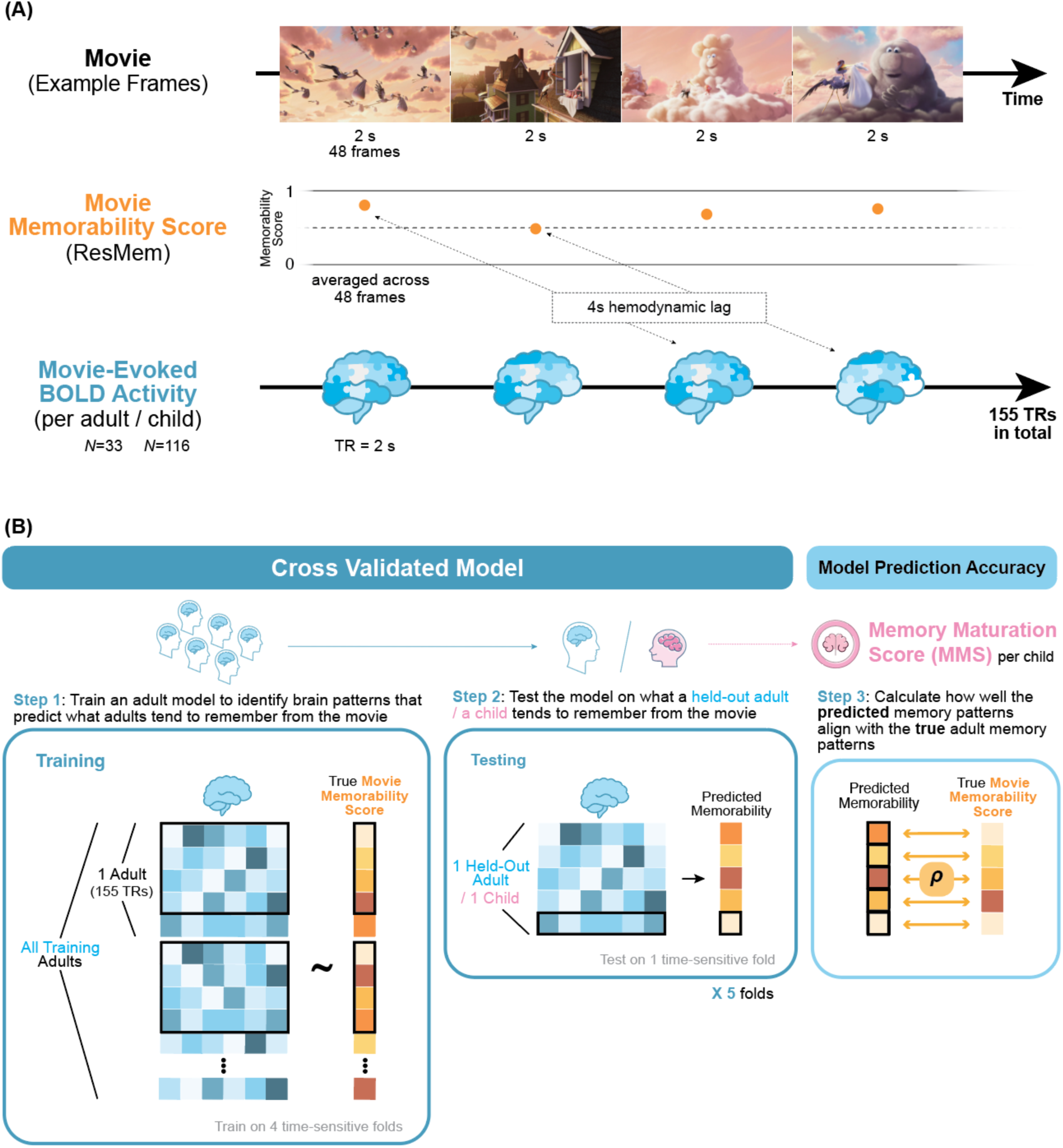
Overview of the Cross-Validated Model. **A.** The movie (*Partly Cloudy*; example frames shown) was segmented into 2-s snippets, each containing 48 frames. The memorability score for each snippet was computed by averaging the frame-wise memorability scores estimated by ResMem and aligned with the movie-evoked BOLD activity for each TR using a 4-s hemodynamic lag. In total, 155 movie memorability scores were aligned with 155 TRs of brain activity. **B.** To construct the cross-validated model, we trained a five-fold Ridge–PCA regression model (see *Methods and Materials* for details) to predict movie memorability from adults’ whole-brain activity and tested the model on either a held-out adult or a child. For adult testing, the model was trained on all adults except the held-out participant. For child testing, the model was trained on all adult participants. Prediction accuracy for each test participant was quantified as the correlation between the model-predicted memorability and the original movie memorability. For children, this prediction accuracy served as the *memory maturation score (MMS)*, with higher values indicating greater similarity between children’s and adults’ neural patterns of memory.

### Creating a Model-Derived Measure of Predicted Memorability Using Movie-Watching Brain Data

The first question is whether brain activity evoked during movie-watching reflects what adults tend to remember. In order to compare children’s brain data to adults’ and calculate a maturation score, we first need to establish that adults’ brain patterns predict one another. Prior work has predicted memorability from adults’ perceptual brain data for static images (18, 20), yet different approaches are required for continuous dynamic stimuli that lack clear stimulus boundaries.

We developed a cross-validated model (Fig. 2B) to predict memorability scores using whole-brain blood oxygenation level-dependent (BOLD) activity. Memorability scores are computed using ResMem, which is a deep neural network trained to predict adult memory performance by assigning each image a value between 0 (consistently forgotten across people) and 1 (consistently remembered; 17). ResMem-predicted memorability has been shown to predict which images adults (11) and 3–5-year-old children (19) tend to remember better than behavioral memory estimates from an independent adult sample, suggesting that ResMem captures stable, stimulus-level differences in memorability. We computed 155 predicted memorability scores—shared across participants— by averaging ResMem predictions across the 48 movie frames associated with each TR (26). The 155 BOLD timepoints were divided into five time-locked folds. A Ridge-PCA cross-validation model was trained on four folds of the whole-brain BOLD data from all but one adult to predict memorability scores and then tested on the held-out adult using the remaining fold (see Fig. 2B and *Methods and Materials* for details). A significant model prediction would indicate that the model captures brain signatures associated with which frames adults tend to remember from the movie, and is able to translate its predictions to both new participants and to new movie frames.

Averaged across the 33 leave-one-out cross-validated models, each tested on a different held-out adult, brain activity during movie watching significantly predicted movie memorability (all results in this study are Fisher-transformed, *M_rho_*=.07, *p*<.001; Fig. 3A). This finding demonstrates that perceptual brain activity contains reliable brain signatures associated with memorability.

**Fig. 3.**
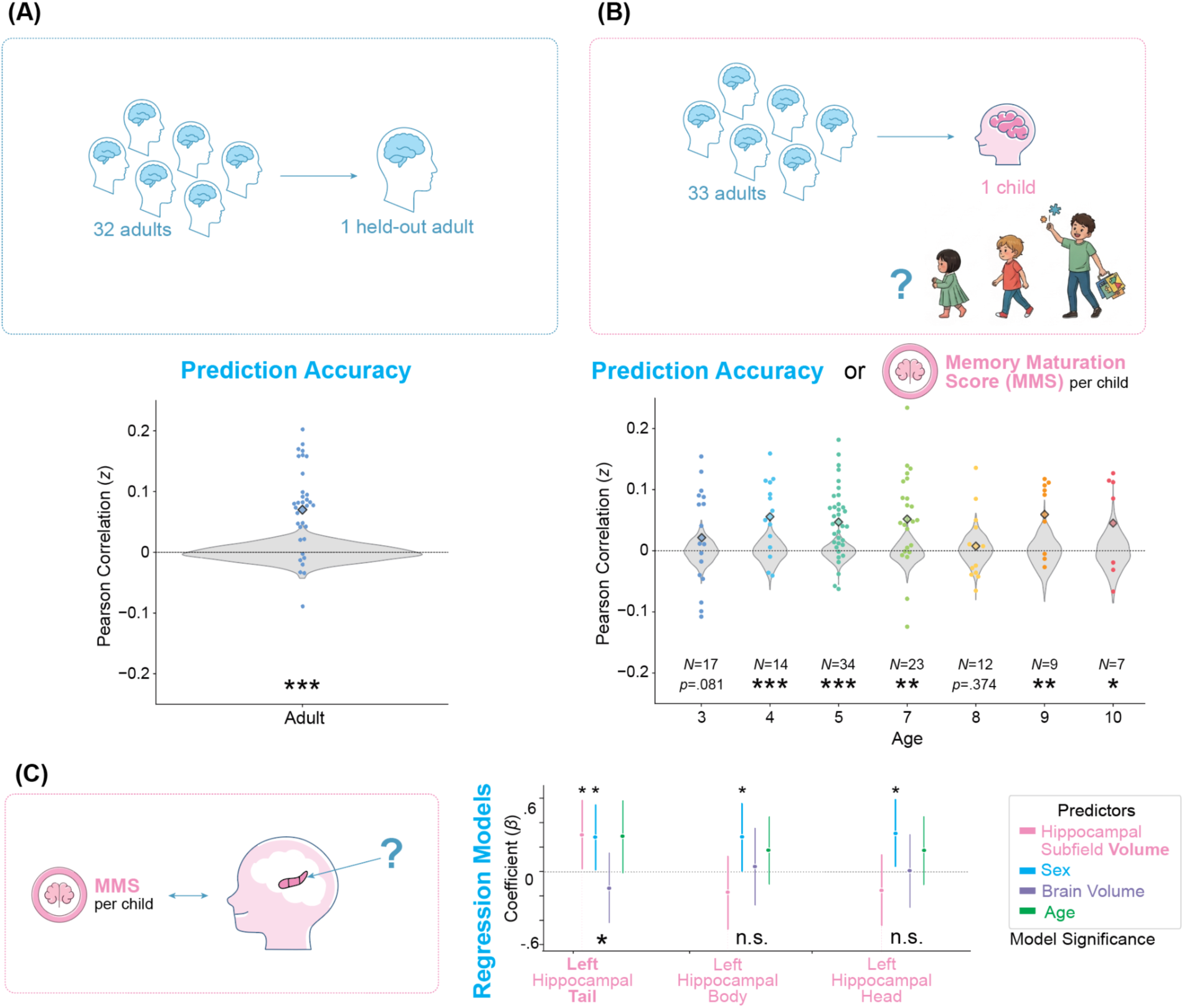
Prediction Accuracy of Memorability (Pearson Correlation) from the Cross-Validated Models Using Whole-Brain Data and Its Relations to Hippocampal Volume. **Top: A.** A model was trained on all but one adult’s perceptual brain data to predict adult behavioral memory patterns (ResMem-predicted memorability scores) and then tested on each held-out adult in a leave-one-subject-out analysis (*N_adult_*=33). **B.** A model was trained on all 33 adults using the same procedure and then tested on each child. Each dot represents a testing participant (adult or child). The colored diamond indicates the mean prediction accuracy. Gray null distributions were created by circular time-shifting memorability scores within each testing fold and recalculating the mean Pearson correlation 1000 times. All correlation results were Fisher-transformed in this study. **Bottom: C.** Linear regression models predicting memory maturation scores (prediction accuracies from **B**) in children aged 3–5 from hippocampal subfield volume (left tail, body, or head), sex, total brain volume, and age. Separate models were fit for each hippocampal subfield. Dots represent standardized regression coefficients (*β*) and vertical lines indicate 95% confidence intervals. Overall model significance is shown below the vertical lines, and significant predictors are indicated above the corresponding vertical lines (*β* estimates). \**p*<.05, \*\**p*<.01, \*\*\**p*<.001.

### Applying the Adult Model to Predict Maturity of Children’s Brain Signatures of Memory

Because adults’ perceptual brain signatures reflect their predicted memorability in the leave-one-out cross-validated model, we trained a cross-validated model on all adults following the same procedures and applied it to each child’s whole-brain data. Using the model trained on adult brain data, successful model prediction of movie memorability from children’s brain data would suggest that children exhibit adult-like brain signatures of memory. Specifically, when a child’s brain data are entered into the cross-validated model, a stronger correlation (i.e., higher prediction accuracy) between the model-predicted behavioral memory patterns and the ResMem-predicted memorability indicates more adult-like sensitivity to memorability in the child’s brain. Accordingly, the model prediction accuracy (mean Pearson correlation coefficient) can be viewed as the child’s *memory maturation score*. This term is used throughout the remainder of this paper.

We applied the cross-validated model trained on all 33 adults to each child and averaged the predicted memory maturation score by age (Fig. 3B). The model successfully predicted mature memory patterns from children’s brain data at ages 4 (*N*=14, *M_rho_*=.06, *p*<.001) and 5 (*N*=34, *M_rho_*=.05, *p*<.001), but not at age 3 (*N*=17, *M_rho_*=.02, *p*=.081). Relative to the model’s theoretical ceiling (i.e., the maximum possible prediction accuracy in the leave-one-out adult analysis, *M_rho_*=.07; see the previous subsection), prediction accuracy reaches 82% and 69% of the theoretical ceiling for ages 4 and 5, respectively. These findings suggest that brain signatures of memory mature around age 4. This replicates prior behavioral findings that mature memory behaviors, as predicted by ResMem, emerge around age 4 (19), while extending this finding to the neural domain and enabling investigation of the neural developmental mechanisms underlying this maturation. Together, these results suggest that age 4 represents a critical developmental window during which children’s memory encoding becomes more mature, as reflected in the maturation of predictable behavioral memory patterns. More broadly, these findings demonstrate the ability of our cross-validated model to predict children’s memory maturation from their perceptual brain data.

In older age groups, children also exhibited mature brain signatures of memory at ages 7 (*N*=23, *M_rho_*=.05, *p*=.001, 76% of theoretical ceiling), 9 (*N*=9, *M_rho_*=.06, *p*=.007, 86% of theoretical ceiling), and 10 (*N*=7, *M_rho_*=.06, *p*=.049, 67% of theoretical ceiling), but not at age 8 (*N*=12, *p*>.05, 11% of theoretical ceiling). The absence of mature memory-related brain signatures in 8-year-olds may reflect a transitional phase during which hippocampal connectivity with the anterior semantic network is reorganized (27) to meet the growing demands of self-deriving knowledge through integrating multiple pieces of memories in school learning (28), resulting in greater variability in brain signatures. However, we refrain from overinterpreting the nonsignificant results at age 8, as the small sample size (*N*=12) may have limited statistical power. In the next subsection exploring neural evidence of the memory shift around age 4, we focus on the 65 children between ages 3 and 5, as the memory shift within this age range has been supported by both neural and behavioral (19) findings.

### Structural Maturation of the Hippocampus Predicts Children’s Memory Maturation

So far, we have identified both neural and behavioral evidence that, relative to age 3, children around age 4 show more mature mnemonic processing and correspondingly more adult-like links to predicted memorability. This robust shift in mnemonic processing signatures coincides with the fading of infantile amnesia, suggesting a possible association between these two phenomena. Because the hippocampus has been implicated both in the structural mechanisms underlying the fading of infantile amnesia and in the functional decoding of the memorability effect, an important next step is to test whether hippocampal structural maturation associates with the maturation of children’s memory brain signatures.

We conducted linear regression analyses to predict children’s memory maturation scores from the volume of each hippocampal subfield among 3-5-year-olds (*N*=65). Because activation patterns in the left hippocampal head, body, and tail have been shown to functionally reflect stimulus memorability in adults (20), we focused on the left hippocampal subfields for this analysis (but report results for the right subfields in *Table S1*). Hippocampal subfield volumes were obtained from manual anatomically-based subfield segmentations across three raters. To determine whether hippocampal volume independently accounted for variance in children’s memory maturation, each regression model also included total brain volume, age (29), and sex (30) as covariates (all predictors were z-scored), given their established associations with hippocampal volume. In total, we fit three linear regression models to data from the 3–5-year-olds (one for each hippocampal subfield), each predicting children’s memory maturation scores from these four predictors (see formula below). Across models, the predictor corresponding to hippocampal subfield volume varied (left head, body, or tail), whereas the remaining covariates were held constant.

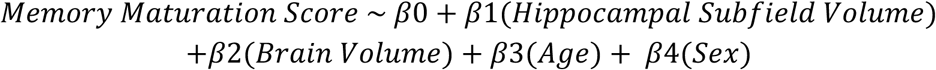

As shown in Fig. 3C, we found that the regression model including left hippocampal tail volume significantly predicted memory maturation (*p*=.02, AIC=168, *adj. R^2^*=.13), with a caveat that this effect became marginally significant after false discovery rate (FDR) correction (*p*=.05, *q*<.05). In this model, both larger left hippocampal tail volume (*β*=.31, *p*=.03, *adj. R^2^*=.059) and male sex (*β*=.28, *p*=.04, *adj. R^2^*=.055) significantly predicted memory maturation score, with the left hippocampal tail volume accounting for more variance (according to the *adj. R^2^*). Brain volume and age were not significant predictors (*p*s > .05). This finding is important for two reasons: (i) the relationship between hippocampal volume growth and memory maturation cannot be explained simply by general increases in brain size; and (ii) the prior behavioral finding that children show mature memory patterns around age 4 likely reflects the development of memory-specific systems involving the hippocampus that occurs alongside with age. By contrast, the models with the left hippocampal head or body volume were not significant (*p*s>.05; see *Table S1 for* detailed statistics and additional models including the right hippocampus).

Together, these regression results suggest that increases in the left hippocampal tail volume, rather than general developmental changes in age or overall brain volume, predominantly account for children’s memory maturation in early childhood. These results further highlight a tight link between the neural mechanisms underlying the fading of infantile amnesia and the maturation of predicted behavioral memory patterns around age 4.

### Developmental Trajectory of Functional Regions that Predict Children’s Memory Maturation

We have shown that structural maturation of the hippocampus predicts children’s memory maturation, indexed by the maturity of their memory brain signatures. However, memory processing is not localized to a single brain region, but instead arises from distributed neural networks across the brain. We therefore expanded our scope by asking what brain regions functionally predict children’s memory maturation at different developmental stages, particularly before and after the memory shift around age 4.

To address this question, we parcellated each participant’s MNI-registered functional data into 224 functional regions using the Brainnetome atlas, including 188 cortical regions and 36 subcortical regions (31, 32). We then conducted a parcel-based searchlight analysis following the same procedures as the whole-brain cross-validated model trained on adults to predict each child’s memory maturation, but restricted the brain data to each parcellated region. In this way, the 224 cross-validated models trained on adults’ regional brain data enabled us to estimate the contribution of each brain region to children’s memory maturation across developmental stages. To increase statistical power, we combined adjacent age groups with relatively small sample sizes (*N_age 3-4_*=31, *N_age 5_*=34, *N_age 7-8_*=35, *N_age 9-10_*=16).

Through visual inspection, unthresholded model prediction accuracies across the 224 parcellated regions (Fig. 4A; see also *Figure S1A* for all brain views) revealed that, with increasing age, children’s brain signatures of memory became increasingly correlated with those of adults, as expected. Upon applying a significance threshold (*p*<.05, FDR-corrected, *q*<.05; Fig. 4B; see also *Figure S1B* for all brain views), we observed a clear neural developmental trajectory of memory across childhood.

**Fig. 4.**
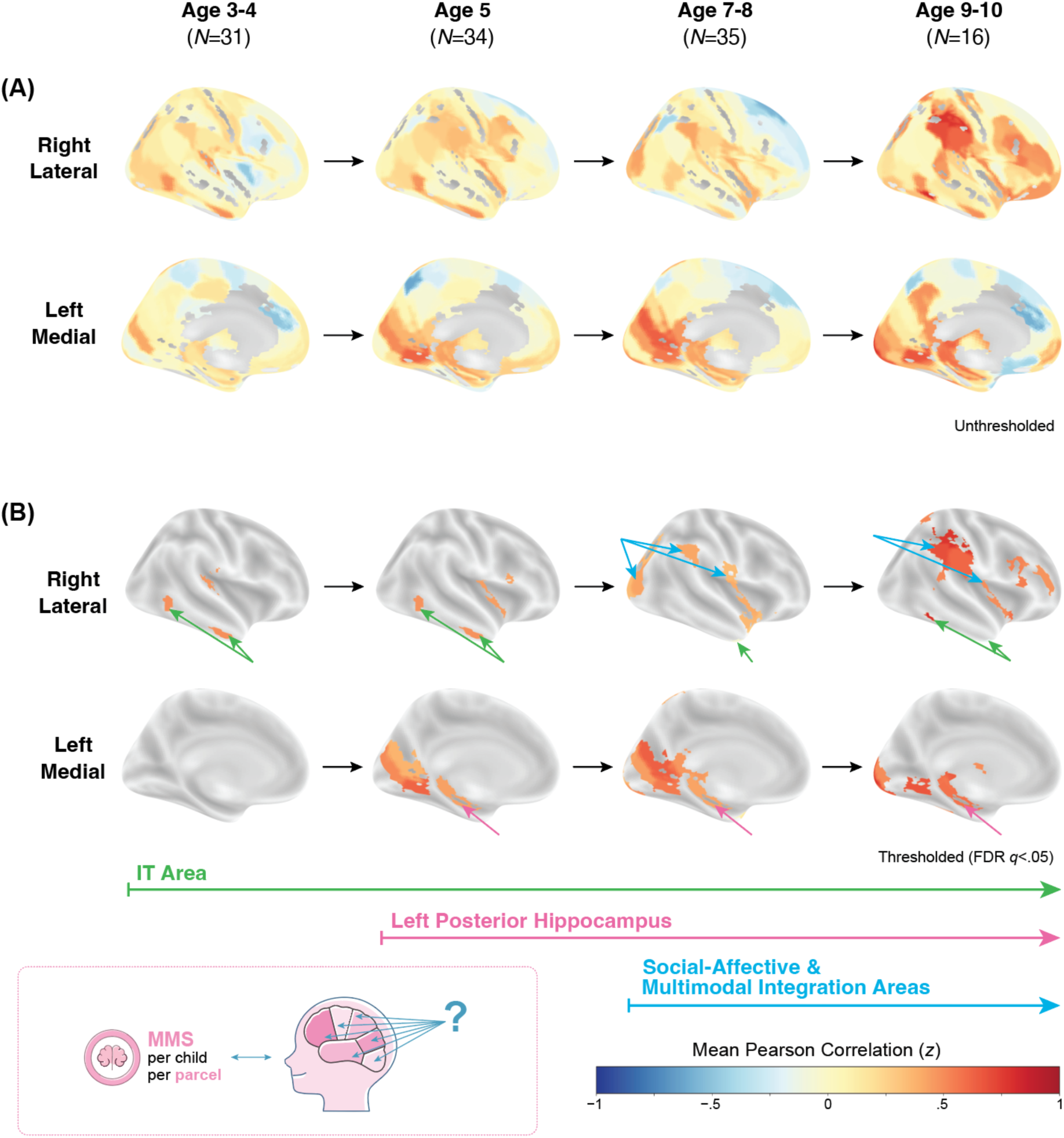
Prediction Accuracy (Pearson Correlation) of the Cross-Validated Models Using Parcellated Brain Data. Functional data from children and adults are parcellated into 224 regions using the Brainnetome atlas (31, 32). For each parcellated region, a cross-validated model was trained on brain data from 33 adults to predict memorability and then tested on children’s brain data from the corresponding region using the same training procedures as in Fig. 2B. **A.** Unthresholded model prediction accuracies across age groups demonstrate increasingly mature brain signatures of memory with age. **B.** Thresholded model prediction accuracies (*p*<.05, FDR-corrected, *q*<.05) reveal a neural developmental trajectory of memory maturation: the inferior temporal gyrus (associated with visual perception) functionally predicts children’s memory maturation by age 3; the left posterior hippocampus (associated with memory processing) by age 5; and regions involved in social-affective processing and multimodal integration by age 7.

Starting at age 3 (and continuing through age 10), the inferotemporal (IT) region, which supports high-level visual processes such as object recognition, significantly predicted children’s memory maturation (see *Table S2* for detailed statistics). Starting at age 5 (and continuing through age 10), brain signatures in the left posterior hippocampus predict children’s memory maturation. This is consistent with the structural findings reported in the previous subsection showing that increases in left hippocampal tail volume from ages 3 to 5 predict children’s memory maturation. Additionally, starting at age 5, regions related to language processing (inferior frontal gyrus), early visual processing (V1 cortex), emotion (cingulate gyrus), and motor control (basal ganglia) also begin to reflect children’s memory maturation. Starting at age 7 (and continuing through age 10), higher-order regions related to social-affective processing (posterior superior temporal sulcus, insular gyrus), multimodal integration (inferior parietal lobule, lateral occipital cortex), attention modulation (thalamus), and somatosensory representation (postcentral gyrus) emerge to significantly predict children’s memory maturation.

Together, these findings demonstrate a developmental trajectory of memory brain signatures, progressing from perceptual to memory-related regions in early childhood and then to higher-order cognitive regions by late childhood. Our cross-validation model paints the following picture: At age 3, children already show mature visual processing in the IT area. At age 5, the emergence of more mature hippocampal function appears to support the observed behavioral memory maturation, although these memories may still lack the richer semantic and social-affective representations supported by higher-order regions that mature around age 7.

## Discussion

Overall, this study delineates the developmental trajectory of distinct neural mechanisms that support adult-like, mature memory. Starting at ages 3-4, children show mature memory processing in the inferotemporal (IT) gyrus, suggesting that high-level visual information (e.g., object category) is already represented in memory at this age. Starting at age 5, functional activity in the left-posterior hippocampus begins to predict children’s memory maturation, suggesting that memory traces at this age contain more semantic information than those at age 3. Notably, we find converging evidence from a separate structural analysis: volume increases in the left-posterior hippocampus from ages 3 to 5, rather than increases in age, predicted children’s memory maturation. This finding complements prior behavioral and our current whole-brain cross-decoding findings showing maturation of memory patterns around age 4 (19), suggesting that this maturation is more closely tied to the age-related development of memory-specific neural systems, including the hippocampus, than to increases in age alone. Starting at age 7, regions involved in social-affective processing and multimodal integration also become predictive of children’s memory maturation, suggesting increasing contributions of richer semantic and integrative processing to children’s memory representations. This resonates with a recent fMRI study showing that the encoding of complex relational structures across distinct items continues to develop between ages 7 and 12 (33). Together, these new findings advance our understanding of early memory development by revealing the developmental timing of distinct neural mechanisms underlying mature memory processing and identifying what children tend to remember from dynamic visual information and what information is likely to persist across development.

### Leveraging ResMem-Predicted Memory Patterns to Predict Memory Maturation

The present findings open new avenues for studying memory development in populations for whom explicit memory measures are difficult or impossible to obtain (e.g., young children and individuals with cognitive decline). By leveraging perceptual brain responses, ResMem and our cross-validated model enable the investigation of memory without requiring explicit memory tests (18, 34). Additionally, our findings suggest that chronological age is a poor proxy for the latent processes underlying memory maturation. In the present study, hippocampal volume predicts memory maturation better than age, underscoring the importance of directly measuring the neural mechanisms that support memory development. Our cross-validated model provides a generalizable framework for studying memory by computationally extracting a memory maturation score from perceptual brain data without relying on assumptions about individual differences or being tied to specific experimental paradigms. Consequently, this approach facilitates more direct comparisons of memory development across studies, tasks, and populations and provides a common metric for characterizing memory maturation across development.

Despite the absence of explicit behavioral memory measures in the current study, converging evidence suggests that ResMem accurately captures children’s memory patterns by around age 4. Prior work shows that stimuli predicted to be more memorable by ResMem are also more likely to be remembered by 4-year-olds, and this relationship is strengthened for older versus younger 4-year-olds (19). The neural findings from the present study further support this relationship, demonstrating that mature memory brain signatures can be decoded from children’s perceptual brain activity by age 4. A potential concern is whether our cross-validated computational framework reflects general visual features or attentional engagement rather than memory-specific representations. However, previous studies demonstrate that (i) ResMem-derived estimates cannot be reduced to visual properties of the images (e.g., color, pattern; 13); (ii) brain signatures of memorability are distinct from those of visual processing in the brain (18); and (iii) memorability and attention are independent predictors of memory (35–37). Together, these findings support the interpretation that our model specifically isolates memory-related processing.

Nevertheless, our goal is not to model all cognitive processes contributing to memory, but rather to develop a sufficiently sensitive model for tracing the maturation of memory patterns within the visual domain. Future research could use ResMem to predict which dynamic visual information is likely to be remembered by young children. Such approaches are particularly valuable because traditional memory measures in young children are often constrained by language abilities (38) and influenced by extraneous factors, including fatigue during lengthy experimental sessions and frustration associated with recall failure. Predicting what children are likely to remember at each age could help educators and caregivers identify memorable learning materials and design learning environments that support retention (39).

### Why Does the Left Hippocampal Tail Predict Memory Maturation in Early Childhood?

Memory researchers have proposed a functional framework of memory along the hippocampal long axis, extending from the posterior hippocampus, which preferentially supports spatial and contextual processing, to the anterior hippocampus, which is more strongly associated with semantic and conceptual processing (see review: 40). Consistent with this framework, fMRI meta-analyses have shown that semantically rich emotion processing in young children is more strongly associated with the anterior hippocampus, whereas memory processing is more strongly associated with the posterior hippocampus (41). The functional dissociation between anterior and posterior hippocampal regions results in the anterior hippocampus, together with its connections to anterior brain regions, developing more gradually than the posterior hippocampus (42), as semantic, conceptual, and emotional processing depend on cross-modal integration and the accumulation of linguistic and cognitive experience. Correspondingly, in the present study, the left posterior hippocampus begins to show mature brain signatures of memory at age 5. Yet, regions involved in multimodal integration and social-affective memory processing, which requires extensive processing (43), do not exhibit mature patterns until approximately age 7, supporting this account of delayed maturation of the coupling between the anterior hippocampus and semantic processing regions.

In addition, previous work has demonstrated functional lateralization of the hippocampus, with the left hippocampus preferentially supporting episodic, contextual, and language-related memory and the right hippocampus preferentially supporting navigation (44). Because stimulus memorability influences episodic memory and is driven in part by semantic content of stimuli (12, 13), it likely depends more strongly on the left hippocampus. Consistent with this interpretation, functional activity in the left hippocampus, but not the right, predicts adult memory patterns (20). Moreover, a study comparing anatomical scans with out-of-scanner memory performance averaged across task paradigms (source memory and spatial location) found that larger left hippocampal tail volume predicts episodic memory performance in children, but not adults, whereas hippocampal head and body volumes predict memory performance in adults, but not children (45). Together, these findings are consistent with the present results showing that the left hippocampal tail’s volume and functional activity predict children’s memory maturation, likely reflecting both the earlier maturation of the posterior relative to the anterior hippocampus and the left-lateralized contribution of episodic and semantic memory processes to stimulus memorability.

### Why Does the Male Sex Show Greater Memory Maturation in Early Childhood?

The posterior and the right hippocampus, which preferentially support spatial processing (40, 44), have been linked to male advantages in spatial cognition. Although spatial processing abilities emerge early in development, male children exhibit relatively advanced spatial abilities by approximately age 4.5 (46–48). This developmental difference suggests that, relative to female children, male children may exhibit more mature memory-related processing in the posterior and the right hippocampus. Consistent with this conjecture, we find that male sex predicts higher memory maturation scores in both models involving the left posterior hippocampus and the right hippocampal subfields, although the overall models involving the right subfields were not significant (see *Table S1*). This suggests that sex is not the central predictor of memory maturation, particularly given the significant model involving the left posterior hippocampus but not the right hippocampal subfields. This is notable because sex has been identified as an important determinant of hippocampal volume growth (29). Yet, our findings suggest that hippocampal volume and function is more directly informative about memory maturation than sex itself. Future research could investigate boundary conditions under which sex predicts memory maturation beyond the contribution of hippocampal function, perhaps using stimuli that place less emphasis on spatial and semantic processing.

### Insights into Infantile Amnesia

The goal of this study was to examine how the specific contents that children tend to remember across development differ from those remembered by adults, around the timepoint when infantile amnesia attenuates, and memory patterns mature. Prior research on infantile amnesia has often focused on whether early memories are *lost* due to poor consolidation (21, 49, 50) or become *inaccessible* due to unsuccessful retrieval (51), but has rarely examined the encoding process, especially the nuanced interactions between child-specific neural maturation and stimulus-specific properties during encoding. Yates and colleagues (23) recently found that infants’ hippocampi reflect the encoding of static images by 1 year old, emphasizing the brain’s ability to encode episodic memory in early infancy. However, the stability of these episodic memory traces remains unknown, especially when presented with more naturalistic, dynamic visual information. The current study suggests that despite the presence of memory encoding in infancy and early childhood, these memory traces are vulnerable representations of the visual information. This vulnerability during encoding in early childhood is constrained by the maturation of key neural regions (i.e., hippocampus and the higher-order processing regions), and the accelerated loss of early memories beyond typical memory decay or interference may reflect immaturity within these key regions.

Specifically, 3-year-old children show less mature memory patterns in both neural activity and memory behavior (19), yet they already show mature brain signatures of memory within the inferotemporal (IT) cortex, suggesting mature representations of high-level visual information such as object categories. This early maturation of the IT cortex can support consistent encoding of high-level visual information in 3-year-olds. Indeed, prior work has shown that 3-year-olds tend to remember and forget the same visual stimuli as one another, although these memory patterns differ from those of adults (19). By contrast, it is not until maturation of the left posterior hippocampus around age 5 and higher-order processing regions around age 7 that children begin to organize perceptual input using richer semantic information in a more adult-like manner. These findings suggest that maturation of these key neural regions determine the maturity of children’s memory representations, and consequently influence the long-term stability or decay of early memories. Identifying these key neural regions provides targeted loci for future investigations into the consolidation and retrieval mechanisms underlying infantile amnesia, as well as the enduring influence of infantile memories despite their apparent inaccessibility later in life. Furthermore, contemporary reviews have highlighted developmental changes in the hippocampus, self-narrative abilities, and memory representational formats as major driving forces behind the attenuation of infantile amnesia (1). The present findings extend this framework by suggesting that maturation of perceptual and higher-order cortical regions also contributes to the developmental trajectory of memory.

### Insights for Future Research and the General Public

This study represents an initial step toward tracing detailed memory characteristics as infantile amnesia attenuates, opening new research directions across disciplines and providing tools with potential applications in both research and education.

For developmental researchers, the present findings identify neural underpinnings of the memory maturation around age 4, enabling more targeted investigations of the neural changes that drive this transition. In addition, our cross-validated model makes it possible to assess children’s memory maturation using perceptual fMRI data without requiring explicit memory measures from young children. More broadly, this approach of training a cross-validated model on healthy adults and using prediction accuracy to quantify how closely each tested participant resembles typical adults could provide an exciting avenue for examining other forms of cognitive divergence in future research.

For memory researchers, these findings suggest that mature mnemonic processing of stimuli during encoding is shaped not only by experience (19), but also by neural maturation of the left posterior hippocampus. Because degeneration of the left temporal lobe is associated with memory patterns that diverge from those of healthy adults in individuals with dementia (52), future work could investigate whether the left posterior hippocampus serves as an early biomarker of cognitive decline and a potential target for intervention through neural stimulation. Likewise, researchers could modify ResMem by increasing the weighting of modules that process visual, relative to semantic, information to determine whether such changes better predict the memory performance of 3-year-olds and younger children.

For medical researchers and practitioners, the memory maturation score derived from our cross-validated model can facilitate the early identification of children who are at risk for later deficits in delayed recall. Prior work has shown that a range of early-life conditions, including maternal anxiety, impaired maternal glycemic control, adoption from international orphanages, and preterm birth, can alter hippocampal development and are associated with subsequent impairments in recall (53). By estimating memory maturation from infants’ perceptual brain responses, our approach may enable the early screening of children at risk for later memory difficulties and inform the development of interventions aimed at promoting typical hippocampal and memory development.

For parents and educators, the present findings suggest that ResMem could be used to select memorable dynamic materials for children as early as age 4. Prior work has shown that memorable stimuli require fewer memory resources, allowing individuals to retain a greater number of items overall (54). Thus, curating more memorable images and videos for learning may help students retain knowledge more efficiently (12, 55).

### Concluding Remarks

By identifying the neural underpinnings that support children’s tendency to remember the same types of information as adults, this study reveals a developmental neural trajectory that may also underlie the attenuation of infantile amnesia. More broadly, the present work uses a cross-decoding framework for tracing the neural signatures of mature memory patterns and provides a tool for assessing divergence from a typical population across other cognitive domains using naturalistic stimuli.

## Methods and Materials

### Extracting fMRI and Memorability Data

We used an existing fMRI dataset (25), in which data were acquired on a 3-Tesla Siemens Tim Trio scanner with custom 32-channel head coils for different ages. The dataset contains separate functional scans (TR=2 s; voxel sizes for different batches: 3.13 mm isotropic for 33 adults, 3 7yo, 20 8-12yo; 3 mm isotropic for the remaining participants; all functional data were subsequently upsampled to 2 mm isotropic voxels in normalized space; adult FOV=256 mm, child FOV=192 mm; see more acquisition details in 25) and T1-weighted structural scans (1 mm isotropic voxels) for each participant (*N_adult_*=33; *N_child_*=116: *N_Age 3_*=17, *N_Age 4_*=14, *N_Age 5_*=34, *N_Age 7_*=23, *N_Age 8_*=12, *N_Age 9_*=9, *N_Age 10_*=7). All scans were preprocessed using SPM8 (56) and custom software written in MATLAB and R. Functional images were realigned to the first image of each run, coregistered to each participant’s anatomical image, and normalized to Montreal Neurological Institute (MNI) space. Functional scans were acquired while participants watched a 5.6-min muted Disney Pixar film, *Partly Cloudy* (24 frames/s). Children aged 11 (*N*=4) and 12 (*N*=2) were excluded because of the small sample sizes. We also excluded the opening Pixar logo and closing credits, restricting our analyses to the main movie segment (5.2 min) and the corresponding 155 TRs of brain data, aligned to the movie with a 4-s hemodynamic lag.

To compute memorability scores for the 155 movie snippets corresponding to the 155 TRs of brain data, we first extracted a memorability score (ranging from 0 for highly forgettable to 1 for highly memorable) for each movie frame using ResMem (17) and then averaged the scores across the 48 frames contained within each TR (2 s). Because prior work has shown that the memorability of a dynamic segment correlates with the average memorability of its constituent moments (26), we used the average frame-level memorability score as a proxy for the likelihood that each 2-s movie segment would be remembered by adults. In total, we extracted a neural time series comprising 155 TRs and a corresponding time series of 155 memorability scores.

ResMem combines the outputs of a deep neural network that uses the architectures of ResNet-152, which captures semantic features of input images, and AlexNet, which captures visual features, and passes these combined features through fully connected layers to generate a predicted memorability score (17). Although ResMem was trained on large image memorability datasets composed of ∼70,000 real-world photographs (e.g., landscapes, people, and pets), it has generalized to predict the memorability of abstract stimuli, such as symbols (10) and artwork (11), as well as abstract, dynamic stimuli, such as co-speech gestures (12). ResMem also successfully predicts children’s memory for scene images (19) and adults’ real-world memory for artworks in a museum better than groups of out-of-sample adults (11)—the very population on which ResMem was trained. Taken together, these findings suggest that ResMem can robustly predict the memorability of a cartoon movie.

### Cross-Validated Model

#### Predicting adult memory patterns from adults’ brain data

As shown in Fig. 2B, we used Ridge regression with Principal Component Analysis (Ridge-PCA) to assess whether movie-evoked brain data (BOLD activity) reflects adults’ predicted behavioral memory patterns of the movie content (ResMem-predicted memorability scores) using leave-one-subject-out cross-validation (*N*=33). In step 1, because there are substantially more voxels (N= 500804; voxels that were neither zero nor empty across all adults were considered) than training participants (*N*=32), we extracted principal components (PCs) across all the TRs of one randomly selected training adult and applied these PCs to all the TRs of the remaining adults in the training set and the participant being tested. The number of PCs equals the number of training participants. In step 2, we divided each training adult’s brain data into five equal folds of timepoints, each containing 31 consecutive movie timepoints. Such time-locked fold partitioning maximizes the differences in movie content between the training and testing sets. We trained the Ridge-PCA model on four folds of movie-evoked brain data from all training adults to predict the ResMem-predicted memorability of the movie contents corresponding to these four folds. Each movie timepoint was associated with a ResMem-predicted score. In step 3, we tested this model on the left-out fold from a held-out participant’s brain data (dimensionality-reduced), yielding predicted memorability scores for that fold. Steps 2 and 3 were repeated five times so that predicted memorability scores were obtained for all five folds of the held-out participants’ brain data. In step 4, we computed Pearson correlations between the predicted and the true memorability scores within each fold and averaged the correlation coefficients across the five folds to obtain the mean prediction accuracy for each held-out adult viewing the held-out scenes. Because there were 32 training adults for each held-out participant and the PCs in step 1 were derived from one randomly selected training adult, steps 1–4 were repeated 32 times, each time selecting PCs from a different training adult. Thus, each held-out adult was predicted by 32 adult models. Model prediction accuracy for each held-out adult is the mean prediction accuracy across these 32 models.

In step 5, we conducted a permutation test using circular time shifting (applying a random shift between 1 and the length of each fold minus 1; 56, 57) to permute the predicted memorability within each fold and recalculate the model prediction accuracy over 1000 iterations to generate a null distribution. Circular time shifting disrupts the within-timepoint correspondence between brain data and memorability scores while preserving temporal dependencies in both the brain activity and the memorability scores, thereby specifically testing the predictability of within-timepoint memorability from brain data. Model significance for each held-out adult was determined as the proportion of null values greater than or equal to the observed mean model prediction accuracy (one-tailed test). Notably, circular time shifting is a conservative method because the permuted memorability series may be shifted relative to itself only by a small offset, such as 1 timepoint, meaning that the null distribution may contain non-null values. As a result, this approach yields a conservative test of significance while maintaining robust control over spurious effects.

#### Predicting adult memory patterns from children’s brain data

We trained a second cross-validated model using the same procedure, except that the training set included all 33 adults and the model was tested on each child. This produces a *memory maturation score* for each child, reflecting the extent to which the child’s memory-related brain signatures can be predicted by a model trained on adults.

### Extracting Hippocampal Subfield Volumes

We first up-sampled the T1-weighted images to a higher resolution using ITK-SNAP (59) to improve segmentation precision. Two trained experimenters (Dice similarity >.8 for all subfields) then manually segmented the hippocampal head, body, and tail from each child’s up-sampled T1-weighted image following existing guidelines (60, 61). A third experimenter visually inspected the segmentations prior to extracting subfield volumes.

### Extracting Brain Volume

Brain volume was obtained via automated brain segmentation using a neural network *deepbet* (62), followed by visual inspection.

### Parcel Searchlight Analysis

Because all functional data were pre-registered to the adult MNI space, we applied the adult Brainnetome atlas (32) to extract BOLD activity from 246 parcels for all participants. The Brainnetome group also provides a child atlas, enabling correspondence between adult and child parcels. The child Brainnetome atlas contains 224 parcels because some parcels are subdivided into multiple parcels in the adult atlas over development (31). Accordingly, we down-sampled the adult atlas to match the 224 parcels in the child atlas (31) by (i) merging 18 adult parcels (in the superior frontal gyrus, insular gyrus, and lateral occipital cortex) into 8 parcels, and (ii) removing 12 adult parcels (related to inferotemporal gyrus, superior parietal lobule, insular gyrus, cingulate gyrus) that lacked corresponding subregion labels in the child atlas (see parcel details in *Table S3*). An experimenter visually inspected parcellation coverage and accuracy for each participant. Before parcellation, we down-sampled the adult Brainnetome atlas (91×109×91) to match the dimensions of the BOLD data (79×95×68) and applied an affine transformation (rotation, scaling, and shearing). Transformation parameters were derived from one randomly selected child and applied to all children, and from one randomly selected adult and applied to all adults. Using parameters from different participants for the affine transformation produced visually equivalent parcellations.

Within each parcel, we followed the same procedure used to train the whole-brain cross-validated model (see subsection *Cross-Validated Model*). Although the number of voxels within each parcel was substantially smaller than that of the whole brain, we still applied PCA to reduce dimensionality, facilitate computation, and maintain a consistent analytical pipeline. Principal components were extracted across voxels within each parcel.

## Acknowledgements

We thank Richardson and colleagues (25) for making their dataset publicly available. We thank Dossay Oryspayev for supporting our use of the high-performance computing resources from the Research Computing Center (RCC) at the University of Chicago. We also thank Yi Fu and Vicky Zhang for their help on manual hippocampal subfield segmentation, and the BLRB group at the University of Chicago for providing insightful comments. Data from this study can be accessed through the Open Science Framework following this link: https://osf.io/e4r7s (code and segmentation data will be made available upon publication of the paper). Once OSF is down, all experimental data and code can be accessed through Zenodo (DOI: 10.5281/zenodo.23928822). Early versions of this work were presented as a conference talk at the Vision Sciences Society (VSS) in 2026 and as a conference poster at FLUX: The Society for Developmental Cognitive Neuroscience in 2025. This work was supported by the National Science Foundation under Grant No. 2441710 and by an Alfred P. Sloan Foundation Fellowship in Neuroscience to WAB.

## Author Contributions

XHG conceptualized the study, analyzed the data, constructed the visualizations, and prepared the first draft. YCL and WAB provided supervision, aided in idea generation and analysis approach, and edited subsequent drafts. All authors reviewed the manuscript prior to submission.

