## Supplemental Materials for "Brain Decoding from a Deep Neural Network of Visual Memory Reveals the Maturation Trajectory of Memory Systems across Ages 3-7"

Xiaohan (Hannah) Guo https://orcid.org/0000-0002-8938-5458

Yuan Chang Leong https://orcid.org/0000-0003-2499-2393

Wilma A. Bainbridge https://orcid.org/0000-0002-7554-0736

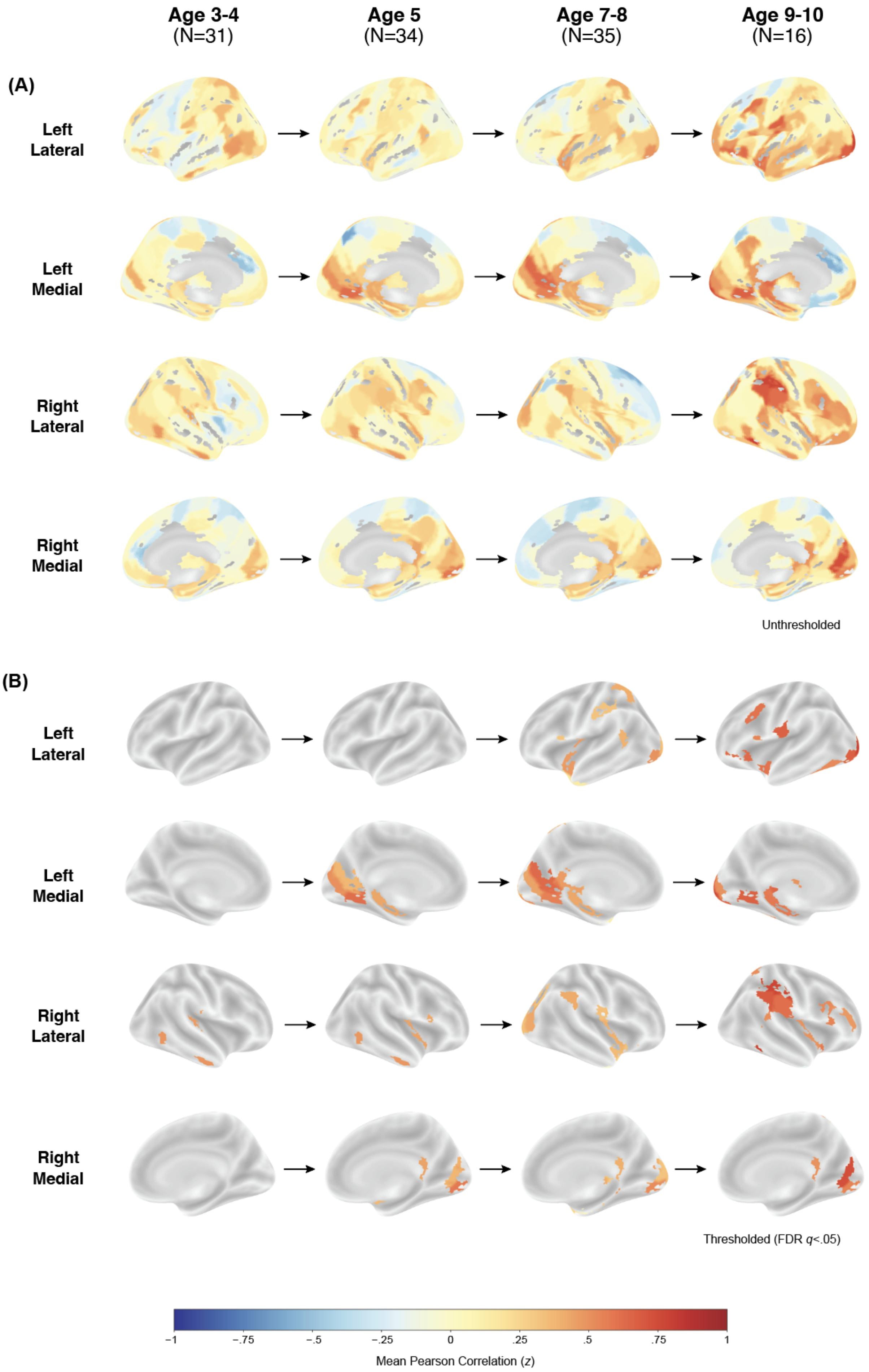
Age 3-4
(N=31)
Age 5
(N=34)
Age 7-8
(N=35)
Age 9-10
(N=16)
(A)
Left Lateral
Left Medial
Right Lateral
Right Medial
Unthresholded
(B)
Left Lateral
Left Medial
Right Lateral
Right Medial
Thresholded (FDR q<.05)
−1
−.75
−.5
−.25
0
.25
.5
.75
1
Mean Pearson Correlation (z)

**Fig. S1. All Brain Views for the Prediction Accuracy (Pearson Correlation) of the Cross-Validated Models Using Parcellated Brain Data. A.** Unthresholded model prediction accuracies across age groups. **B.** Thresholded model prediction accuracies (*p*<.05, FDR-corrected, *q*<.05) across age groups.

**Table S1. Detailed Statistics for the Structural Analysis Models Including Hippocampal Subfield Volume.**

*Significant p-values are bolded.*

| y | | Predictors | | | | | | | | | | |
|---|---|---|---|---|---|---|---|---|---|---|---|---|
| | | Hippocampal Subfield Volume | | Brain Volume | | Age | | Sex | | | | |
| | | *β* | *p* | *β* | *p* | *β* | *p* | *β* | *p* | AIC | *p* | Adj. $R^2$ |
| Memory Maturation Score | Left Head | -.15 | .30 | .006 | .97 | .17 | .22 | .32 | .02 | 171 | .08 | .07 |
| | Left Body | -.17 | .25 | .04 | .80 | .17 | .21 | .28 | .05 | 171 | .07 | .08 |
| | Left Tail | .31 | **.03** | -.13 | .37 | .29 | .06 | .28 | **.04** | 168 | **.02** | .13 |
| | Right Head | -.14 | .36 | .03 | .87 | .16 | .25 | .31 | **.03** | 172 | .09 | .07 |
| | Right Body | .13 | .36 | -.07 | .63 | .09 | .52 | .32 | **.02** | 172 | .09 | .07 |
| | Right Tail | -.12 | .40 | -.007 | .96 | .09 | .54 | .31 | **.03** | 172 | .09 | .07 |

**Table S2. Prediction Accuracies (Memory Maturation) for Parcellated Brain Regions That Functionally Predict Adult-Like Memory Processing in Each Age Group.**

*M = mean correlation (r); p = permutation p-value. "—" indicates the parcel was not significant for that age group. Age in years; N = number of children per group.*

| Related Function | Region | Parcel | Age Group | | | | | | | |
|---|---|---|---|---|---|---|---|---|---|---|
| | | | 3-4 (N=31) | | 5 (N=34) | | 7-8 (N=35) | | 9-10 (N=16) | |
| | | | M | p | M | p | M | p | M | p |
| Object recognition, high-level vision | Inferior Temporal Gyrus | **A37elv_R**, extreme lateroventral area37 | — | — | — | — | — | — | .08 | *<.001* |
| | | **A20r_L**, rostral area 20 | — | — | — | — | .01 | *<.001* | — | — |
| | | **A20r_R**, rostral area 20 | — | — | — | — | -.01 | *<.001* | -.02 | *<.001* |
| | | **A20il_R**, intermediate lateral area 20 | .05 | *<.001* | .05 | *<.001* | — | — | — | — |
| | Superior Temporal Gyrus | **A38m_L**, medial area 38 | — | — | — | — | .02 | *<.001* | — | — |
| | | **A38m_R**, medial area 38 | — | — | — | — | .03 | *<.001* | — | — |
| | | **A41/42_R**, area 41/42 | .05 | *<.001* | — | — | — | — | — | — |
| | | **A38l_L**, lateral area 38 | — | — | — | — | .05 | *<.001* | .06 | *<.001* |
| | | **A38l_R**, lateral area 38 | — | — | — | — | .04 | *.002* | — | — |
| Episodic memory, spatial / contextual / associative memory | **Hippocampus** | **cHipp_L**, caudal hippocampus | — | — | .04 | *.001* | .04 | *.001* | .06 | *.001* |
| Emotion processing | **Amygdala** | **mAmyg_R**, medial amygdala | — | — | .04 | *.002* | — | — | — | — |
| Motor control, reward-based learning, procedural learning | **Basal Ganglia** | **vmPu_L**, ventromedial putamen | — | — | — | — | — | — | .05 | *.004* |
| | | **vmPu_R**, ventromedial putamen | — | — | .04 | *.001* | — | — | — | — |
| | | **dlPu_R**, dorsolateral putamen | — | — | — | — | .03 | *.007* | — | — |
| Language processing | **Inferior Frontal Gyrus** | **A44d_R**, dorsal area 44 | — | — | — | — | — | — | .05 | *.006* |
| | | **A45c_R**, caudal area 45 | — | — | — | — | — | — | .05 | *.006* |
| | | **A44v_L**, ventral area 44 | — | — | — | — | .03 | *.008* | .05 | *.005* |
| | | **A44v_R**, ventral area 44 | — | — | .04 | *.001* | — | — | — | — |
| Auditory processing | **Cingulate Gyrus** | **A23v_L**, ventral area 23 | — | — | — | — | .05 | *<.001* | — | — |
| | | **A23v_R**, ventral area 23 | — | — | .04 | *.001* | .04 | *<.001* | .05 | *.004* |
| Visual processing | **Medio Ventral Occipital Cortex** | **cLinG_R**, caudal lingual gyrus | — | — | .06 | *<.001* | .05 | *<.001* | .05 | *.003* |
| | | **rCunG_L**, rostral cuneus gyrus | — | — | .05 | *<.001* | .04 | *.001* | — | — |
| | | **rCunG_R**, rostral cuneus gyrus | — | — | .04 | *.002* | — | — | .07 | *<.001* |
| | | **cCunG_L**, caudal cuneus gyrus | — | — | — | — | — | — | .05 | *.003* |
| | | **cCunG_R**, caudal cuneus gyrus | — | — | — | — | .03 | *.008* | — | — |
| | | **rLinG_L**, rostral lingual gyrus | — | — | .06 | *<.001* | .06 | *<.001* | .07 | *<.001* |
| | | **vmPOS_L**, ventromedial parietooccipital sulcus | — | — | .04 | *.002* | .06 | *<.001* | — | — |
| Social processing, multimodal integration | **Posterior Superior Temporal Sulcus** | **rpSTS_L**, rostroposterior superior temporal sulcus | — | — | — | — | .04 | *.002* | — | — |
| | | **rpSTS_R**, rostroposterior superior temporal sulcus | — | — | — | — | — | — | .05 | *.003* |
| Attention, multimodal integration | **Inferior Parietal Lobule** | **A39rd_L**, rostrodorsal area 39(Hip3) | — | — | — | — | .03 | *.003* | — | — |
| | | **A39rd_R**, rostrodorsal area 39(Hip3) | — | — | — | — | — | — | .08 | *<.001* |
| | | **A40rd_R**, rostrodorsal area 40(PFt) | — | — | — | — | .04 | *.002* | .07 | *<.001* |
| | | **A39rv_R**, rostroventral area 39(PGa) | — | — | — | — | — | — | .06 | *.002* |

| Related Function | Region | Parcel | Age Group | | | | | | | |
|---|---|---|---|---|---|---|---|---|---|---|
| | | | 3-4 (N=31) | | 5 (N=34) | | 7-8 (N=35) | | 9-10 (N=16) | |
| | | | M | p | M | p | M | p | M | p |
| Affective processing, regulation | **Insular Gyrus** | **vIa_R**, ventral agranular insula | — | — | — | — | .03 | *.007* | .06 | *.004* |
| Object integration, attention | **Lateral Occipital Cortex** | **mOccG_R**, middle occipital gyrus | — | — | — | — | .04 | *<.001* | — | — |
| | | **OPC_L**, occipital polar cortex | — | — | — | — | .03 | *.005* | .08 | *<.001* |
| | | **OPC_R**, occipital polar cortex | — | — | — | — | .03 | *.002* | — | — |
| | | **iOccG_L**, inferior occipital gyrus | — | — | — | — | .05 | *<.001* | .07 | *.001* |
| Visual attention modulation | **Thalamus** | **PPtha_L**, posterior parietal thalamus | — | — | — | — | .04 | *.003* | — | — |
| | | **PPtha_R**, posterior parietal thalamus | — | — | — | — | .04 | *<.001* | — | — |
| | | **Otha_L**, occipital thalamus | — | — | — | — | .05 | *<.001* | .06 | *<.001* |
| | | **Otha_R**, occipital thalamus | — | — | — | — | .03 | *.008* | — | — |
| Visuospatial processing, memory encoding | **Parahippocampal Gyrus** | **A35/36r_R**, rostral area 35/36 | — | — | — | — | .03 | *.007* | — | — |
| | | **A28/34_L**, area 28/34 (EC, entorhinal cortex) | — | — | — | — | .001 | *<.001* | — | — |
| | | **A28/34_R**, area 28/34 (EC, entorhinal cortex) | — | — | — | — | .02 | *<.001* | — | — |
| | | **TI_R**, area TI(temporal agranular insular cortex) | — | — | — | — | .05 | *.001* | — | — |
| Visual processing, attention | **Superior Parietal Lobule** | **A5l_L**, lateral area 5 | — | — | — | — | .04 | *<.001* | — | — |
| High-level visual processing | **Fusiform Gyrus** | **A37mv_L**, medioventral area37 | — | — | — | — | — | — | .05 | *.002* |
| | | **A37lv_L**, lateroventral area37 | — | — | — | — | -.01 | *<.001* | — | — |
| | | **A37lv_R**, lateroventral area37 | — | — | — | — | .001 | *<.001* | — | — |
| Somatosensory representation | **Postcentral Gyrus** | **A1/2/3ulhf_L**, area 1/2/3(upper limb, head and face region) | — | — | — | — | — | — | .07 | *<.001* |
| | | **A1/2/3ulhf_R**, area 1/2/3(upper limb, head and face region) | — | — | — | — | .03 | *.002* | — | — |
| Executive functions | **Middle Frontal Gyrus** | **IFJ_L**, inferior frontal junction | — | — | — | — | — | — | .06 | *<.001* |
| | | **A9/46v_R**, ventral area 9/46 | — | — | — | — | — | — | .05 | *.004* |
| | **Orbital Gyrus** | **A12/47l_L**, lateral area 12/47 | — | — | — | — | — | — | .06 | *.001* |
| Motor execution | **Precentral Gyrus** | **A4t_L**, area 4(trunk region) | — | — | — | — | — | — | -.02 | *<.001* |

### Table S3. Matching of the Adult (1) and Child (2) Brainnetome Atlas Parcel Number.

*L = left hemisphere; R = Right hemisphere*

| Lobe | Gyrus | Anatomical and Modified Cyto-Architectonic Descriptions | Adult Parcel Number | | Child Parcel Number | |
|---|---|---|---|---|---|---|
| | | | L | R | L | R |
| Frontal Lobe | SFG, Superior Frontal Gyrus | A8m, medial area 8 | 1 | 2 | 1 | 2 |
| | | A8dl, dorsolateral area 8 | 3 | 4 | 3 | 4 |
| | | A9m/A9l, medial area 9/lateral area 9 | 5,11 | 6,12 | 5 | 6 |
| | | A6dl, dorsolateral area 6 | 7 | 8 | 7 | 8 |
| | | A6m, medial area 6 | 9 | 10 | 9 | 10 |
| | | A10m, medial area 10 | 13 | 14 | 11 | 12 |
| | MFG, Middle Frontal Gyrus | A9/46d, dorsal area 9/46 | 15 | 16 | 13 | 14 |
| | | IFJ, inferior frontal junction | 17 | 18 | 15 | 16 |
| | | A46, area 46 | 19 | 20 | 17 | 18 |
| | | A9/46v, ventral area 9/46 | 21 | 22 | 19 | 20 |
| | | A8vl, ventrolateral area 8 | 23 | 24 | 21 | 22 |
| | | A6vl, ventrolateral area 6 | 25 | 26 | 23 | 24 |
| | | A10l, lateral area10 | 27 | 28 | 25 | 26 |
| | IFG, Inferior Frontal Gyrus | A44d, dorsal area 44 | 29 | 30 | 27 | 28 |
| | | IFS, inferior frontal sulcus | 31 | 32 | 29 | 30 |
| | | A45c, caudal area 45 | 33 | 34 | 31 | 32 |
| | | A45r, rostral area 45 | 35 | 36 | 33 | 34 |
| | | A44op, opercular area 44 | 37 | 38 | 35 | 36 |
| | | A44v, ventral area 44 | 39 | 40 | 37 | 38 |
| | OrG, Orbital Gyrus | A14m, medial area 14 | 41 | 42 | 39 | 40 |
| | | A12/47o, orbital area 12/47 | 43 | 44 | 41 | 42 |
| | | A11l, lateral area 11 | 45 | 46 | 43 | 44 |
| | | A11m, medial area 11 | 47 | 48 | 45 | 46 |
| | | A13, area 13 | 49 | 50 | 47 | 48 |
| | | A12/47l, lateral area 12/47 | 51 | 52 | 49 | 50 |
| | PrG, Precentral Gyrus | A4hf, area 4(head and face region) | 53 | 54 | 51 | 52 |
| | | A6cdl, caudal dorsolateral area 6 | 55 | 56 | 53 | 54 |
| | | A4ul, area 4(upper limb region) | 57 | 58 | 55 | 56 |
| | | A4t, area 4(trunk region) | 59 | 60 | 57 | 58 |
| | | A4tl, area 4(tongue and larynx region) | 61 | 62 | 59 | 60 |
| | | A6cvl, caudal ventrolateral area 6 | 63 | 64 | 61 | 62 |
| | PCL, Paracentral Lobule | A1/2/3ll, area1/2/3 (lower limb region) | 65 | 66 | 63 | 64 |
| | | A4ll, area 4, (lower limb region) | 67 | 68 | 65 | 66 |
| Temporal Lobe | STG, Superior Temporal Gyrus | A38m, medial area 38 | 69 | 70 | 67 | 68 |
| | | A41/42, area 41/42 | 71 | 72 | 69 | 70 |
| | | TE1.0 and TE1.2 | 73 | 74 | 71 | 72 |
| | | A22c, caudal area 22 | 75 | 76 | 73 | 74 |
| | | A38l, lateral area 38 | 77 | 78 | 75 | 76 |
| | | A22r, rostral area 22 | 79 | 80 | 77 | 78 |
| | MTG, Middle Temporal Gyrus | A21c, caudal area 21 | 81 | 82 | 79 | 80 |
| | | A21r, rostral area 21 | 83 | 84 | 81 | 82 |
| | | A37dl, dorsolateral area37 | 85 | 86 | 83 | 84 |
| | | aSTS, anterior superior temporal sulcus | 87 | 88 | 85 | 86 |
| | ITG, Inferior Temporal Gyrus | A37elv, extreme lateroventral area37 | 91 | 92 | 87 | 88 |
| | | A20r, rostral area 20 | 93 | 94 | 89 | 90 |
| | | A20il, intermediate lateral area 20 | 95 | 96 | 91 | 92 |
| | | A20cl, caudolateral of area 20 | 99 | 100 | 93 | 94 |
| | | A20cv, caudoventral of area 20 | 101 | 102 | 95 | 96 |
| | FuG, Fusiform Gyrus | A20rv, rostroventral area 20 | 103 | 104 | 97 | 98 |
| | | A37mv, medioventral area37 | 105 | 106 | 99 | 100 |
| | | A37lv, lateroventral area37 | 107 | 108 | 101 | 102 |

| Lobe | Gyrus | Anatomical and Modified Cyto-Architectonic Descriptions | Adult Parcel Number | | Child Parcel Number | |
|---|---|---|---|---|---|---|
| | | | L | R | L | R |
| | PhG, Parahippocampal Gyrus | A35/36r, rostral area 35/36 | 109 | 110 | 103 | 104 |
| | | A35/36c, caudal area 35/36 | 111 | 112 | 105 | 106 |
| | | TL, area TL (lateral PPHC, posterior parahippocampal gyrus) | 113 | 114 | 107 | 108 |
| | | A28/34, area 28/34 (EC, entorhinal cortex) | 115 | 116 | 109 | 110 |
| | | TI, area TI(temporal agranular insular cortex) | 117 | 118 | 111 | 112 |
| | | TH, area TH (medial PPHC) | 119 | 120 | 113 | 114 |
| | pSTS, posterior Superior Temporal Sulcus | rpSTS, rostroposterior superior temporal sulcus | 121 | 122 | 115 | 116 |
| | | cpSTS, caudoposterior superior temporal sulcus | 123 | 124 | 117 | 118 |
| Parietal Lobe | SPL, Superior Parietal Lobule | A7c, caudal area 7 | 127 | 128 | 119 | 120 |
| | | A5l, lateral area 5 | 129 | 130 | 121 | 122 |
| | | A7pc, postcentral area 7 | 131 | 132 | 123 | 124 |
| | IPL, Inferior Parietal Lobule | A39c, caudal area 39(PGp) | 135 | 136 | 125 | 126 |
| | | A39rd, rostrodorsal area 39(Hip3) | 137 | 138 | 127 | 128 |
| | | A40rd, rostrodorsal area 40(PFt) | 139 | 140 | 129 | 130 |
| | | A40c, caudal area 40(PFm) | 141 | 142 | 131 | 132 |
| | | A39rv, rostroventral area 39(PGa) | 143 | 144 | 133 | 134 |
| | | A40rv, rostroventral area 40(PFop) | 145 | 146 | 135 | 136 |
| | Pcun, Precuneus | A7m, medial area 7(PEp) | 147 | 148 | 137 | 138 |
| | | A5m, medial area 5(PEm) | 149 | 150 | 139 | 140 |
| | | dmPOS, dorsomedial parietooccipital sulcus(PEr) | 151 | 152 | 141 | 142 |
| | | A31, area 31 (Lc1) | 153 | 154 | 143 | 144 |
| | PoG, Postcentral Gyrus | A1/2/3ulhf, area 1/2/3(upper limb, head and face region) | 155 | 156 | 145 | 146 |
| | | A1/2/3tonla, area 1/2/3(tongue and larynx region) | 157 | 158 | 147 | 148 |
| | | A2, area 2 | 159 | 160 | 149 | 150 |
| | | A1/2/3tru, area1/2/3(trunk region) | 161 | 162 | 151 | 152 |
| Insular Lobe | INS, Insular Gyrus | vIa, ventral agranular insula | 165 | 166 | 153 | 154 |
| | | G/vId/vIg/dIg, hypergranular insula/ventral dysgranular and granular insula/dorsal granular insula | 163,169,171 | 164,170,172 | 155 | 156 |
| | | dIa/dId, dorsal agranular insula/dorsal dysgranular insula | 167,173 | 168,174 | 157 | 158 |
| Limbic Lobe | CG, Cingulate Gyrus | A23d, dorsal area 23 | 175 | 176 | 159 | 160 |
| | | A32p, pregenual area 32 | 179 | 180 | 161 | 162 |
| | | A23v, ventral area 23 | 181 | 182 | 163 | 164 |
| | | A23c, caudal area 23 | 185 | 186 | 165 | 166 |
| | | A32sg, subgenual area 32 | 187 | 188 | 167 | 168 |
| Occipital Lobe | MVOcC, MedioVentral Occipital Cortex | cLinG, caudal lingual gyrus | 189 | 190 | 169 | 170 |
| | | rCunG, rostral cuneus gyrus | 191 | 192 | 171 | 172 |
| | | cCunG, caudal cuneus gyrus | 193 | 194 | 173 | 174 |
| | | rLinG, rostral lingual gyrus | 195 | 196 | 175 | 176 |
| | | vmPOS, ventromedial parietooccipital sulcus | 197 | 198 | 177 | 178 |
| | LOcC, lateral Occipital Cortex | mOccG, middle occipital gyrus | 199 | 200 | 179 | 180 |
| | | V5/MT+, area V5/MT+ | 201 | 202 | 181 | 182 |
| | | OPC, occipital polar cortex | 203 | 204 | 183 | 184 |
| | | iOccG, inferior occipital gyrus | 205 | 206 | 185 | 186 |
| | | msOccG/lsOccG, medial superior occipital gyrus/lateral superior occipital gyrus | 207,209 | 208,210 | 187 | 188 |
| Subcortical Nuclei | Amyg, Amygdala | mAmyg, medial amygdala | 211 | 212 | 189 | 190 |
| | | lAmyg, lateral amygdala | 213 | 214 | 191 | 192 |

| Lobe | Gyrus | Anatomical and Modified Cyto-Architectonic Descriptions | Adult Parcel Number | | Child Parcel Number | |
|---|---|---|---|---|---|---|
| | | | L | R | L | R |
| | **Hipp, Hippocampus** | rHipp, rostral hippocampus | 215 | 216 | 193 | 194 |
| | | cHipp, caudal hippocampus | 217 | 218 | 195 | 196 |
| | **BG, Basal Ganglia** | vCa, ventral caudate | 219 | 220 | 197 | 198 |
| | | GP, globus pallidus | 221 | 222 | 199 | 200 |
| | | NAC, nucleus accumbens | 223 | 224 | 201 | 202 |
| | | vmPu, ventromedial putamen | 225 | 226 | 203 | 204 |
| | | dCa, dorsal caudate | 227 | 228 | 205 | 206 |
| | | dlPu, dorsolateral putamen | 229 | 230 | 207 | 208 |
| | **Tha, Thalamus** | mPFtha, medial pre-frontal thalamus | 231 | 232 | 209 | 210 |
| | | mPMtha, pre-motor thalamus | 233 | 234 | 211 | 212 |
| | | Stha, sensory thalamus | 235 | 236 | 213 | 214 |
| | | rTtha, rostral temporal thalamus | 237 | 238 | 215 | 216 |
| | | PPtha, posterior parietal thalamus | 239 | 240 | 217 | 218 |
| | | Otha, occipital thalamus | 241 | 242 | 219 | 220 |
| | | cTtha, caudal temporal thalamus | 243 | 244 | 221 | 222 |
| | | lPFtha, 3lateral pre-frontal thalamus | 245 | 246 | 223 | 224 |

***SI References***


1. L. Fan, *et al.*, The human brainnetome atlas: a new brain atlas based on connectional architecture. *Cereb. Cortex* **26**, 3508–3526 (2016).

2. W. Li, *et al.*, Brainnetome atlas of preadolescent children based on anatomical connectivity profiles. *Cereb. Cortex* **33**, 5264–5275 (2023).
